## Supplementary FIgures for "Activation loop phosphorylation and cGMP saturation of PKG regulate egress of malaria parasites"

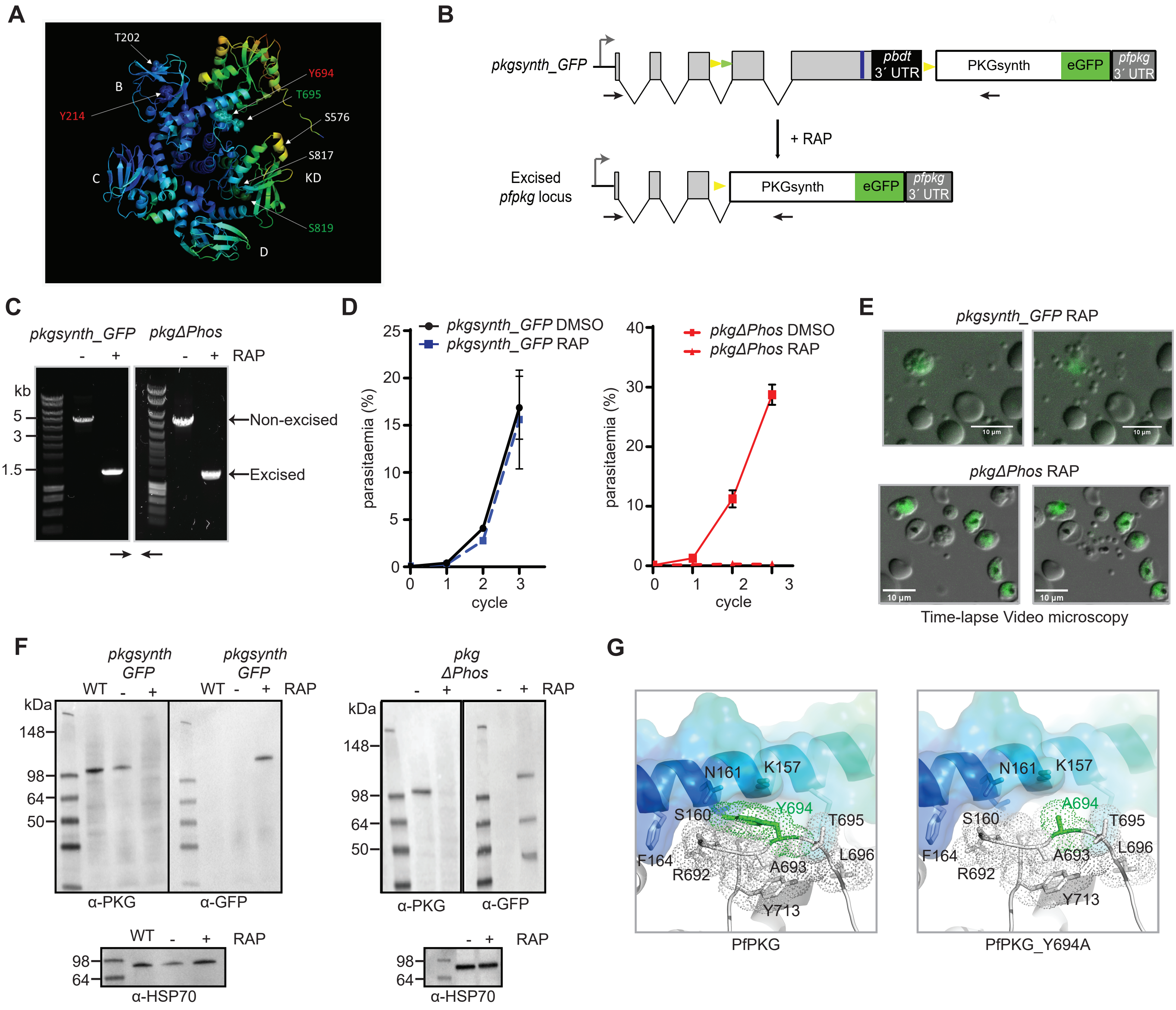

A
T202
B
Y694
T695
Y214
S576
S817
C
KD
S819
D
B
pkgsynth_GFP
pbdt 3´ UTR
PKGsynth
eGFP
pfpkg 3´ UTR
+ RAP
Excised pfpkg locus
C
pkgsynth_GFP
pkgΔPhos
RAP
kb
5
3
1.5
Non-excised
Excised
D
pkgsynth_GFP DMSO
pkgsynth_GFP RAP
parasitaemia (%)
cycle
pkgΔPhos DMSO
pkgΔPhos RAP
E
pkgsynth_GFP RAP
pkgΔPhos RAP
10 µm
Time-lapse Video microscopy
F
pkgsynth GFP
WT
RAP
kDa
148
98
64
50
α-PKG
α-GFP
pkg ΔPhos
α-HSP70
G
N161
K157
S160
Y694
T695
F164
R692
A693
L696
Y713
A694
PfPKG
PfPKG_Y694A

**Fig S1: Phosphosite mutations destabilise the pentagonal architecture of PfPKG**

**(A)** Cartoon representation of the *P. falciparum* PKG x-ray crystal structure (PDB ID: 5DYK) in its apo form where predicted phosphosites are indicated and shown as sticks within colour matching spheres. The cartoon is coloured according to the temperature factor values (B-factors ($Å^2$), spectrum from dark blue for low to red for high). (**B**) Schematic representation of the approach used to create line *pkgsynth_GFP*. Blue line indicates the position targeted by the gRNA. RAP-induced DiCre activity switches expression from wt PKG to a gene replacement with a partially synthetic *pfpkg* gene fused to eGFP. Black arrows; oligonucleotides used for excision PCR. **(C)** Excision PCR showing generation of products after activation of DiCre upon RAP treatment in line *pkgsynth_GFP* and *pkgΔPhos*. Expected sizes of the amplicons corresponding to non-excised and excised fragments are 5kb and 1.4 kb respectively. **(D)** Growth curves showing replication of DMSO-treated (control) or RAP-treated *pkgsynth_GFP* and *pkgΔPhos* lines. Mean values are shown. Error bars: ± SD (n=3). **(E)** Representative still images of a 30 min time-lapse video of mock-treated (grey) or RAP-treated (green) parasites of lines *pkgsynth_GFP* and *pkgΔPhos*. *pkgsynth_GFP* RAP-treated schizonts underwent egress (upper panel), whilst *pkgΔPhos* RAP-treated parasites did not (lower panel). Scale bar, 10 µM. **(F)** (Left panel) Western blot showing expression of PKG (expected MW: 98 kDa) in DMSO-treated *pkgsynth_GFP* schizonts relevant to the parental line (WT). Upon RAP-treatment there is appearance of a signal corresponding to the PKG_GFP fusion (expected molecular weight: 125 kDa). (Right panel) Representative Western blot of *pkgΔPhos* schizonts after DMSO or RAP treatment. Note the multiple bands appearing at the RAP-treated sample. Cytoplasmic HSP-70 was used as a loading control.

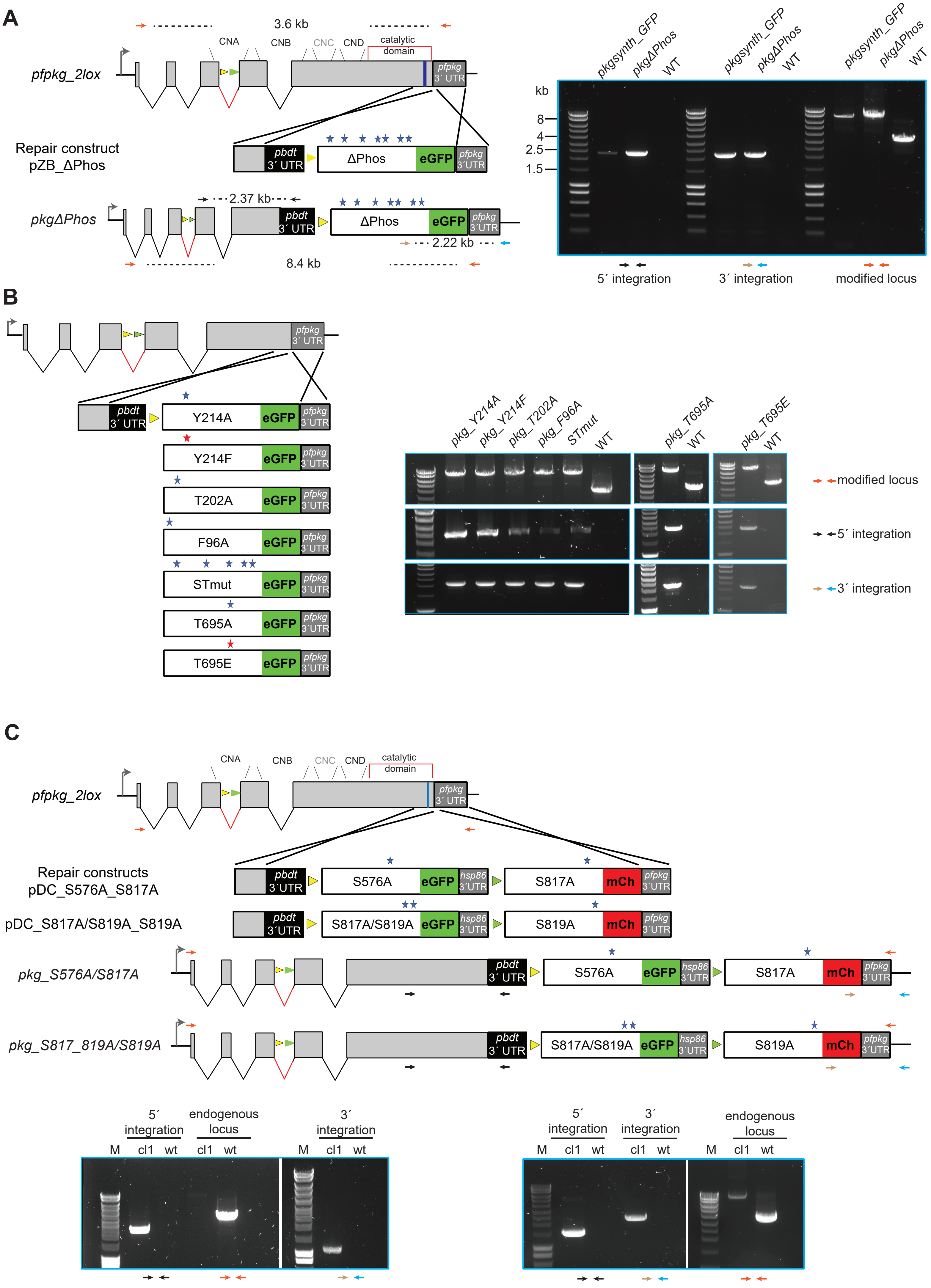
A
3.6 kb
CNA
CNB
CNC
CND
catalytic domain
pfpkg_2lox
pfpkg 3´ UTR
Repair construct pZB_ΔPhos
pbdt 3´ UTR
ΔPhos
eGFP
pkgΔPhos
2.37 kb
2.22 kb
8.4 kb
kb
8
4
2.5
1.5
pkgsynth_GFP
pkgΔPhos
WT
5´ integration
3´ integration
modified locus
B
Y214A
Y214F
T202A
F96A
STmut
T695A
T695E
pkg_Y214A
pkg_Y214F
pkg_T202A
pkg_F96A
STmut
pkg_T695A
pkg_T695E
modified locus
5´ integration
3´ integration
C
Repair constructs pDC_S576A_S817A
pDC_S817A/S819A_S819A
S576A
S817A
S817A/S819A
S819A
mCh
hsp86 3´UTR
pkg_S576A/S817A
pkg_S817_819A/S819A
5´ integration
endogenous locus
3´ integration
M
cl1
wt

**Fig S2: Generation and genotyping of transgenic *P. falciparum* lines**

**(A-B)** Modification strategies and genotyping data for generation of the parasite lines *pkgΔPhos*, *pkgY214A*, *pkgY214F*, *pkgT202A*, *pkgF96A*, *pkgSTmut*, *pkgT695A* and *pkgT695E* respectively. Blue line indicates the position targeted by the gRNA. Relative positions of mutations are depicted by coloured stars. Positions of oligonucleotides used for genotyping by diagnostic PCR are indicated (coloured arrows), and agarose gel electrophoresis of corresponding PCR products are shown. Positions of *lox* sites are indicated with coloured arrowheads (yellow, *loxN*; green, *lox2272*). **(C)** Modification strategy for lines *pkgS576A_S817A* and *pkgS817A/S819A_S819A*. Blue line indicates the position targeted by the gRNA. Relative positions of mutations are depicted by coloured stars. Positions of oligonucleotides used for genotyping by diagnostic PCR are indicated (coloured arrows), and agarose gel electrophoresis of corresponding PCR products are shown. Positions of *lox* sites are indicated with coloured arrowheads (yellow, *loxN*; green, *lox2272*).

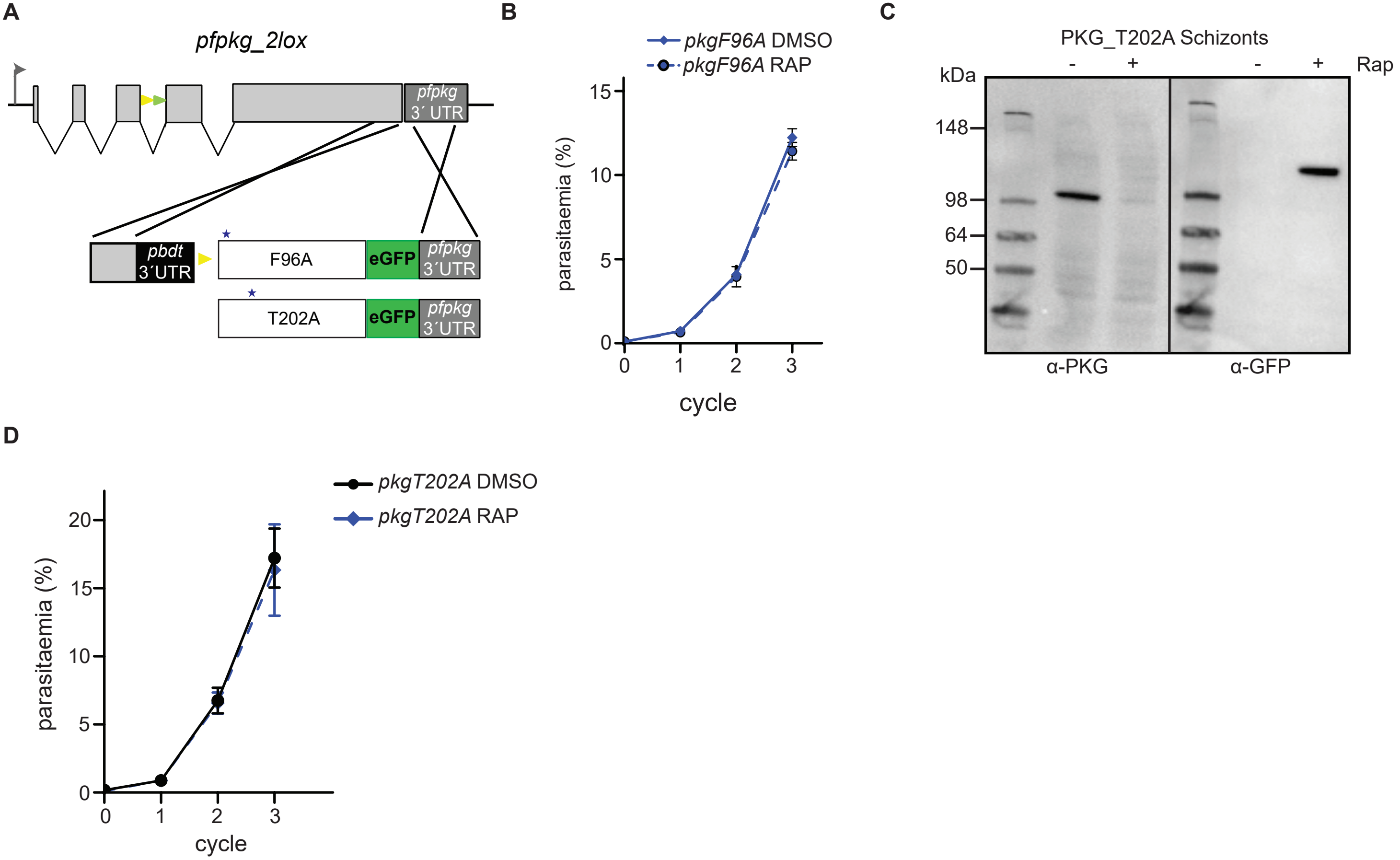
A
pfpkg_2lox
pfpkg
3´ UTR
pbdt
3´UTR
F96A
eGFP
pfpkg
3´UTR
T202A
eGFP
pfpkg
3´UTR
B
pkgF96A DMSO
pkgF96A RAP
parasitaemia (%)
15
10
5
0
0
1
2
3
cycle
C
PKG_T202A Schizonts
-
+
-
+
Rap
kDa
148
98
64
50
α-PKG
α-GFP
D
pkgT202A DMSO
pkgT202A RAP
parasitaemia (%)
20
15
10
5
0
0
1
2
3
cycle

**Fig S3: F96 and T202 mutations do not affect PfPKG function *in vivo*.**

**(A)** Schematic representation of the approach used to create lines *pkgF96A* and *pkgT202A*. Relative positions of the mutated amino acids are denoted by blue stars. **(B)** Growth curve showing replication of DMSO-treated (control) or RAP-treated *pkgF96A*. Mean values are shown. Error bars: ± SD (n=3). **(C)** Western blot showing expression of endogenous PKG in DMSO-treated *pkgT202A* schizonts and of the mutated PKG (T202A) fused to GFP upon RAP-treatment. **(D)** Growth curve showing replication of DMSO-treated (control) or RAP-treated *pkgT202A*. Mean values are shown. Error bars: ± SD (n=3).

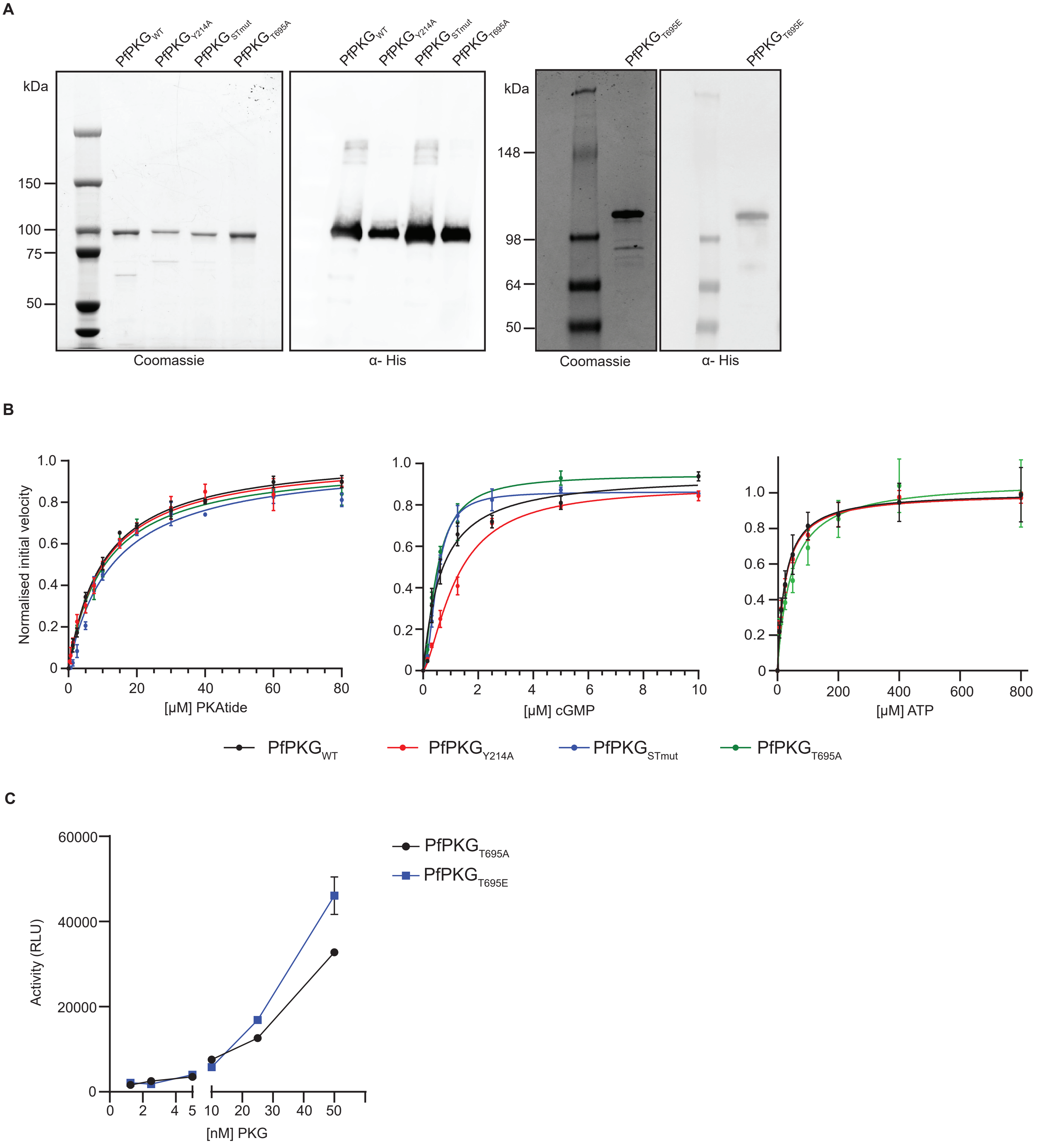

A
PfPKG WT
PfPKG Y214A
PfPKG STmut
PfPKG T695A
PfPKG T695E
kDa
150
100
75
50
148
98
64
Coomassie
α- His
B
Normalised initial velocity
[μM] PKAtide
[μM] cGMP
[μM] ATP
PfPKG WT
PfPKG Y214A
PfPKG STmut
PfPKG T695A
C
Activity (RLU)
60000
40000
20000
[nM] PKG
PfPKG T695A
PfPKG T695E

**Fig S4: Recombinant expression and kinetic properties of the different PfPKG mutants**

**(A)** Coomassie stained gels and western blot analysis of the 5 different recombinant PfPKGs used in this study. **(B)** Determination of PKAtide $K_m$, cGMP $K_A$ and ATP $K_m$ for $PfPKG_{WT}$, $PfPKG_{Y214A}$, $PfPKG_{STmut}$, $PfPKG_{T695A}$. **(C)** $PfPKG_{T695A}$ and $PfPKG_{T695E}$ activity assay showing phosphorylation of PKAtide.

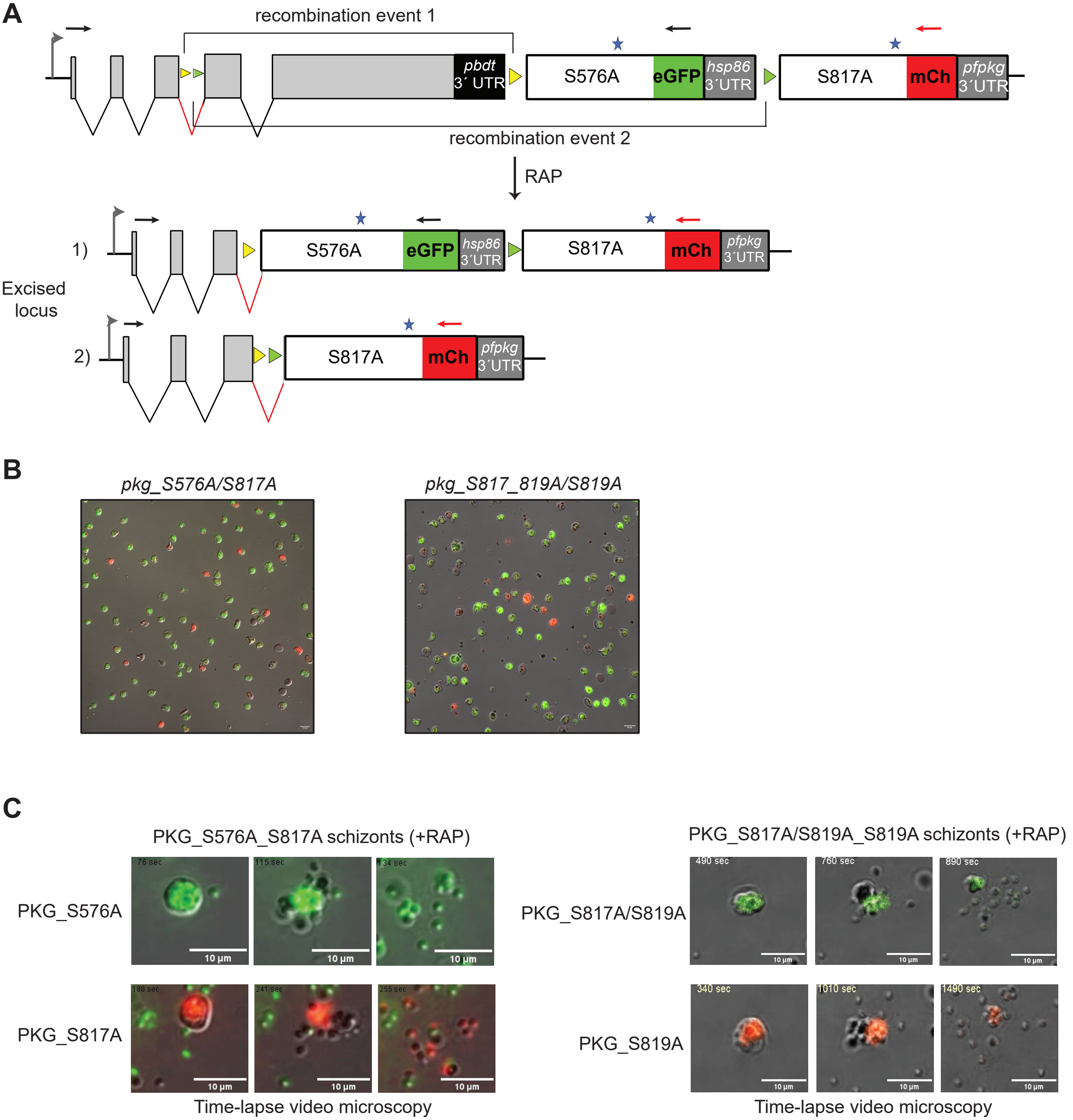

A
recombination event 1
pbdt 3´ UTR
S576A
eGFP
hsp86 3´UTR
S817A
mCh
pfpkg 3´UTR
recombination event 2
RAP
Excised locus
1)
2)
B
pkg_S576A/S817A
pkg_S817_819A/S819A
C
PKG_S576A_S817A schizonts (+RAP)
PKG_S576A
PKG_S817A
76 sec
115 sec
134 sec
188 sec
241 sec
255 sec
10 µm
Time-lapse video microscopy
PKG_S817A/S819A_S819A schizonts (+RAP)
PKG_S817A/S819A
PKG_S819A
490 sec
760 sec
890 sec
340 sec
1010 sec
1490 sec

**Fig S5**: **phosphomutants of the N and C-lobe do not affect parasite viability**

**(A)** Schematic of line *pkgS576A_S817A*. RAP-induced DiCre activity switches expression from WT PKG to either a gene replacement with a S576A mutant fused to eGFP (recombination event 1; PKG_S576A) or to expression of a S817A mutant fused to mCherry (recombination event 2; PKG_S817A). Similar allelic replacement strategy applies to line *pkgS817A/S819A_S819A*. Black and red arrows; oligonucleotides used for identification of both events by diagnostic PCR. **(B)** Representative images from DIC/fluorescence microscopic examination of RAP-treated *pkgS576A_S817A* and RAP-treated *pkgS817A/S819A_S819A* parasites (end of cycle 0), showing both GFP- and mCherry-positive schizonts. Scale bar, 10 μm. **(C)** Stills from time-lapse DIC/fluorescence microscopy of isolated, RAP-treated *pkgS576A_S817A* and *pkgS817A/S819A_S819A* schizonts, showing that all four mutants can undergo egress. Scale bars, 10μm.

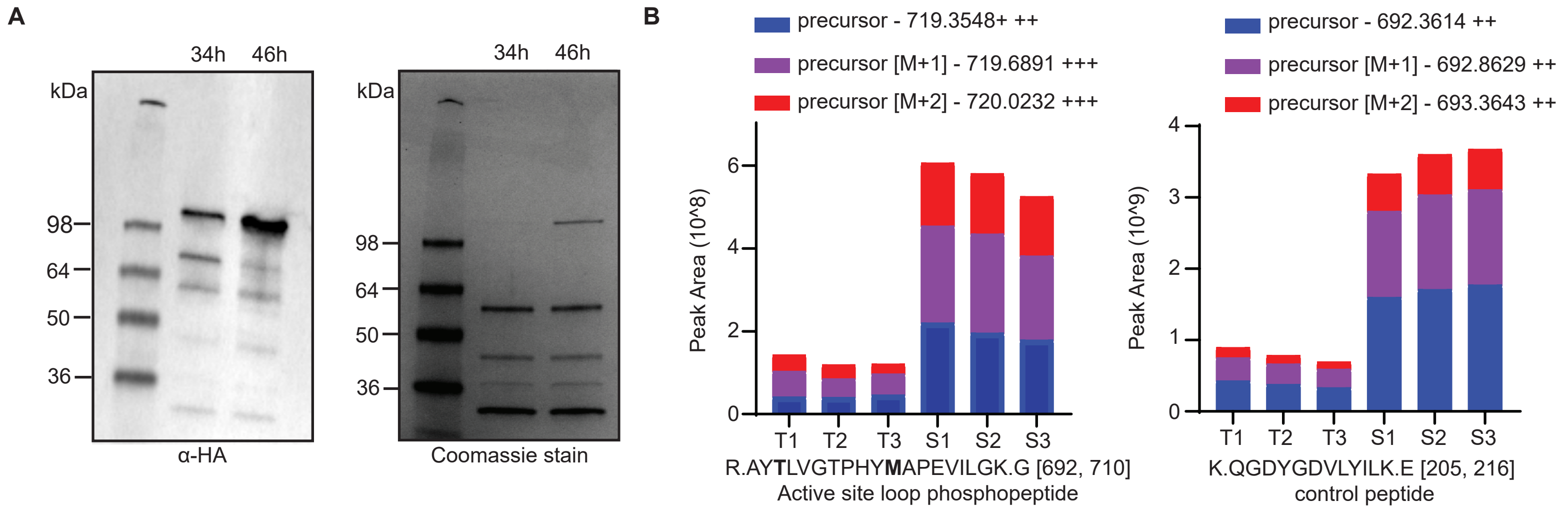
A
34h 46h
kDa
98
64
50
36
α-HA
34h 46h
kDa
98
64
50
36
Coomassie stain
B
precursor - 719.3548+ ++
precursor [M+1] - 719.6891 +++
precursor [M+2] - 720.0232 +++
Peak Area (10^8)
0
2
4
6
T1 T2 T3 S1 S2 S3
R.AY**T**LVGTPHY**M**APEVILGK.G [692, 710]
Active site loop phosphopeptide
precursor - 692.3614 ++
precursor [M+1] - 692.8629 ++
precursor [M+2] - 693.3643 ++
Peak Area (10^9)
0
1
2
3
4
T1 T2 T3 S1 S2 S3
K.QGDYGDVLYILK.E [205, 216]
control peptide

**Fig S6**: **T695 is phosphorylated from trophozoite stages, prior to PKG activation.** **(A)** Western blot and Coomassie stained gel after PKG pulldown showing presence of the protein in both trophozoites (34 h) and schizonts (46 h). **(B)** Peak areas of precursor ion chromatograms are extracted after MS1 filtering for peaks picked based on MS2 peptide identification in three technical replicates at both trophozoite (T1-T3) and schizont samples (S1_S3). Peaks for the phosphorylated T695 peptide were found in all experiments (left panel) showing similar ratio between trophozoite and schizont samples as the control peptide (right panel).

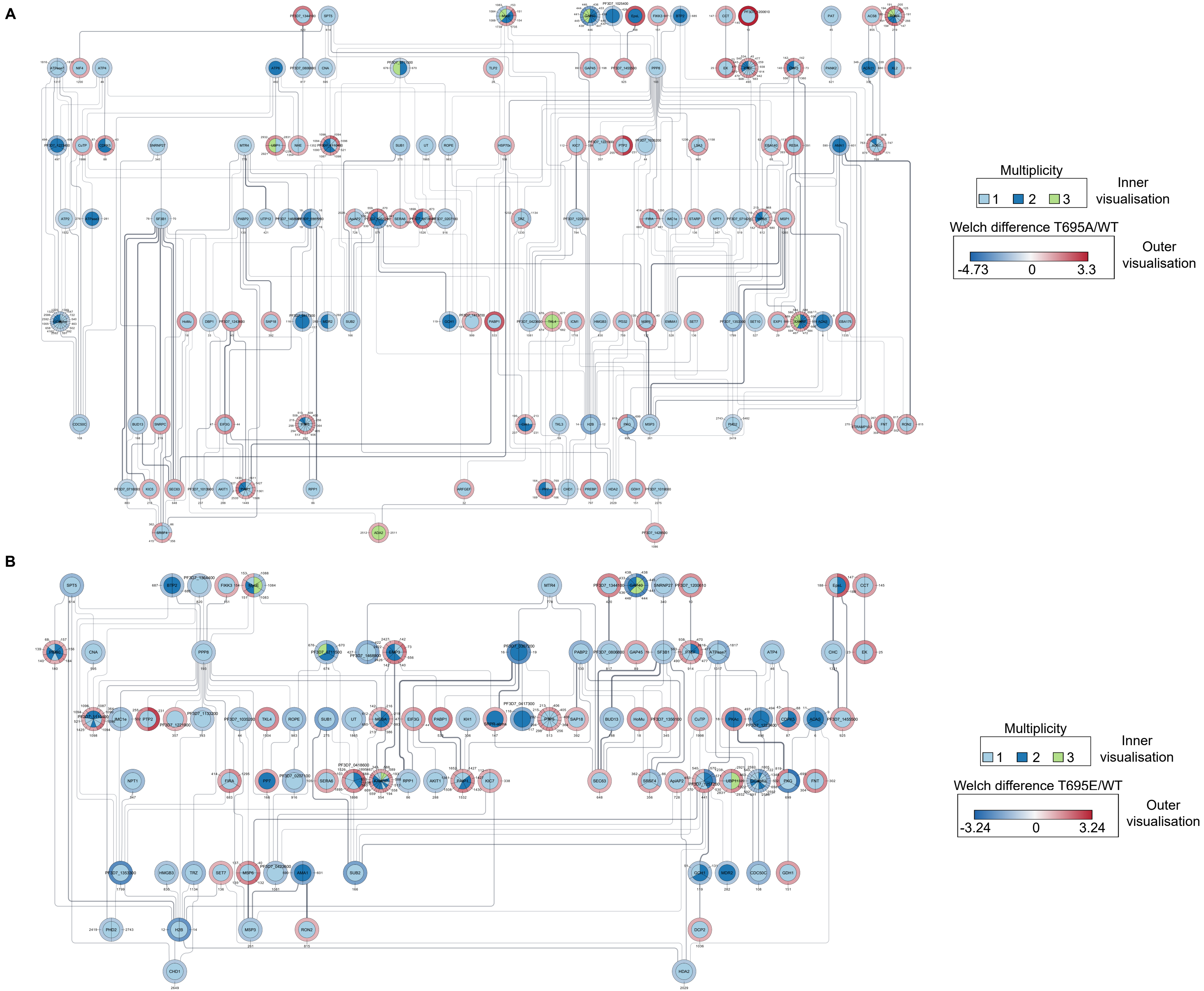
A
Multiplicity
1 2 3
Inner visualisation
Welch difference T695A/WT
-4.73 0 3.3
Outer visualisation
B
Multiplicity
1 2 3
Inner visualisation
Welch difference T695E/WT
-3.24 0 3.24
Outer visualisation

**Fig S7: Protein-protein interaction networks of T695A and T695E deregulated proteins**

**(A)** STRING network (y Files Hierarchical layout) visualizing functional interactions (edges) between proteins (nodes) significantly hypo and hyper-phosphorylated in the T695A mutant over WT. Default STRING clustering confidence score cutoff of 0.4 was used to determine whether two nodes were functionally related. Single proteins are excluded from the schematic. Significantly deregulated phosphosites are depicted on each protein and were coloured according to the Welch difference (outer visualisation) and the multiplicity (number of phosphorylation events in the peptide – inner visualisation). Names or PlasmoDB IDs are displayed on each protein.
**(B)** STRING network (y Files Hierarchical layout) visualizing functional interactions (edges) between proteins (nodes) significantly hypo and hyper-phosphorylated in the T695E mutant over WT. Same clustering parameters and visualisation were used as in (A).

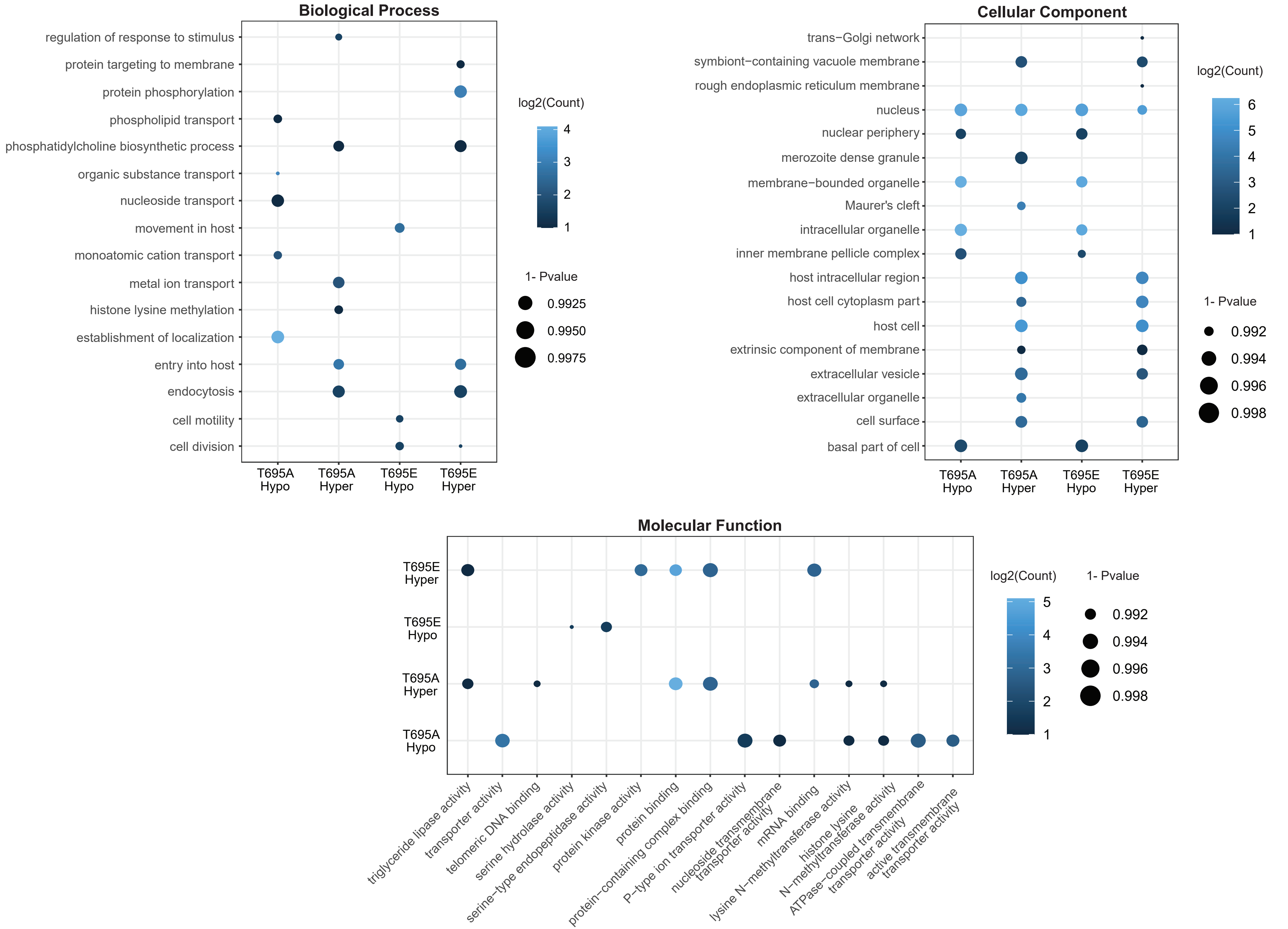
Biological Process
regulation of response to stimulus
protein targeting to membrane
protein phosphorylation
phospholipid transport
phosphatidylcholine biosynthetic process
organic substance transport
nucleoside transport
movement in host
monoatomic cation transport
metal ion transport
histone lysine methylation
establishment of localization
entry into host
endocytosis
cell motility
cell division
T695A Hypo
T695A Hyper
T695E Hypo
T695E Hyper
log2(Count)
4
3
2
1
1- Pvalue
0.9925
0.9950
0.9975
Cellular Component
trans−Golgi network
symbiont−containing vacuole membrane
rough endoplasmic reticulum membrane
nucleus
nuclear periphery
merozoite dense granule
membrane−bounded organelle
Maurer's cleft
intracellular organelle
inner membrane pellicle complex
host intracellular region
host cell cytoplasm part
host cell
extrinsic component of membrane
extracellular vesicle
extracellular organelle
cell surface
basal part of cell
T695A Hypo
T695A Hyper
T695E Hypo
T695E Hyper
log2(Count)
6
5
4
3
2
1
1- Pvalue
0.992
0.994
0.996
0.998
Molecular Function
T695E Hyper
T695E Hypo
T695A Hyper
T695A Hypo
triglyceride lipase activity
transporter activity
telomeric DNA binding
serine hydrolase activity
serine−type endopeptidase activity
protein kinase activity
protein binding
protein−containing complex binding
P−type ion transporter activity
nucleoside transmembrane transporter activity
mRNA binding
lysine N−methyltransferase activity
histone lysine N−methyltransferase activity
ATPase−coupled transmembrane transporter activity
active transmembrane transporter activity
log2(Count)
5
4
3
2
1
1- Pvalue
0.992
0.994
0.996
0.998

**Fig S8: Gene ontology analysis of T695A and T695E phosphoproteomes**

Gene ontology (GO) enrichment analysis of significantly hyper or hypo phosphorylated proteins in T695A and T695E schizonts. GO terms were obtained from the PlasmoDB database. Size of the bubble indicates the level of significance (1-p value) of the enriched GO term and colour density indicates the number of differentially expressed proteins (log2 protein count) associated with the GO term.

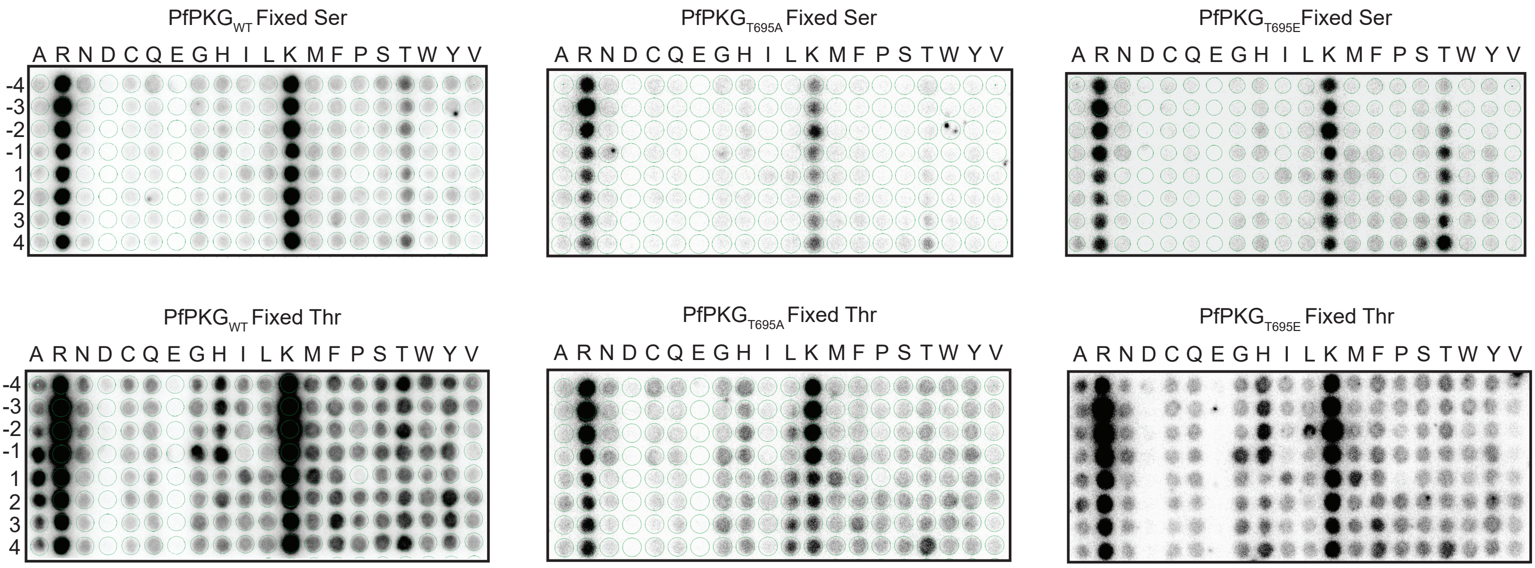
PfPKG$_{WT}$ Fixed Ser
PfPKG$_{T695A}$ Fixed Ser
PfPKG$_{T695E}$ Fixed Ser
PfPKG$_{WT}$ Fixed Thr
PfPKG$_{T695A}$ Fixed Thr
PfPKG$_{T695E}$ Fixed Thr
A R N D C Q E G H I L K M F P S T W Y V
-4
-3
-2
-1
1
2
3
4

**Fig S9: OPAL activity assays and determined sequence logos.**

Autoradiographs of the oriented peptide array libraries showing phosphorylation activity of PfPKGWT, PfPKGT695A and PfPKGT695E (n=1). Library was based on the design A-X-X-X-X-S-X-X-X-X-A or A-X-X-X-X-T-X-X-X-X-A.
